# IL-18 inhibition enlarges lesions, necrotic cores and thickens fibrous caps in *Jak2^V617F^* clonal hematopoiesis-driven atherosclerosis

**DOI:** 10.1101/2025.06.03.657754

**Authors:** Mojdeh Tavallaie, Cheng-Chieh Hsu, Brian D. Hardaway, Huijuan Dou, Trevor Fidler, Eunyoung Kim, Sandra E. Abramowicz, David Ngai, Tong Xiao, Nan Wang, Marit Westerterp, Alan R. Tall

## Abstract

**Background:** Inflammasome activation promotes atherosclerosis in clonal hematopoiesis (CH). Active inflammasomes secrete both IL-1β and IL-18. Plasma IL-18 levels are elevated in *Jak2^VF^*CH. Genetic deficiency of IL-18 has been shown to reduce atherosclerosis in non-CH murine models. However, whether IL-18 inhibition promotes atherosclerosis in control or *Jak2^VF^* CH is unknown.

**Methods and results:** *Ldlr^−/−^* mice were transplanted with bone marrow (BM) from *Mx1-cre Jak2^VF^* (20%) and wild-type (80%) mice or with control BM, fed a Western-type diet (WTD) for 8, 10 or 16 weeks and administered control or IL-18 IgG from 4 weeks onwards.

IL-18 antibody treatment increased plaque collagen content and cap thickness. Unexpectedly, IL-18 antibody treatment increased the size of early lesions and promoted formation of advanced lesions with large necrotic cores in *Jak2^VF^* CH mice. IL-18 antibody treatment was associated with diminished interferon (IFN)-γ and AIM2 levels and reduced macrophage pyroptosis especially in *Jak2^VF^* CH mice. However, IL-18 antibodies increased cleaved Caspase-3 and TUNEL^+^ macrophages (indicating increased apoptosis) and reduced efferocytosis. Sc-RNA-seq analysis showed that IL-18 antibody treatment reduced expression of *MHC class II* genes, a marker of IFN-γ signaling, and of genes mediating efferocytosis (*Mertk and Axl)*, in resident-like macrophage subpopulations in *Jak2^VF^* CH mice. Consistently, IFN-γ injection increased *Axl* and *Mertk* expression in resident peritoneal macrophages.

**Conclusions:** Despite improvements in collagen and fibrous cap thickness in *Jak2^VF^* CH mice, IL-18 antibody treatment increased advanced necrotic lesions, reflecting a shift from pyroptotic to apoptotic cell death coupled with defective efferocytosis, events which were coordinated by reduced IFN-γ signaling. These findings indicate a mixed atherosclerosis phenotype resulting from IL-18 inhibition, advocating for alternative therapeutic strategies.

Inhibition of IL-18 has been considered as a novel therapeutic approach to reduce atherosclerosis and stabilize atherosclerotic plaques. We show that IL-18 antibodies have adverse effects on atherosclerotic lesional necrosis, calling this approach into question.

**Graphical Abstract:** 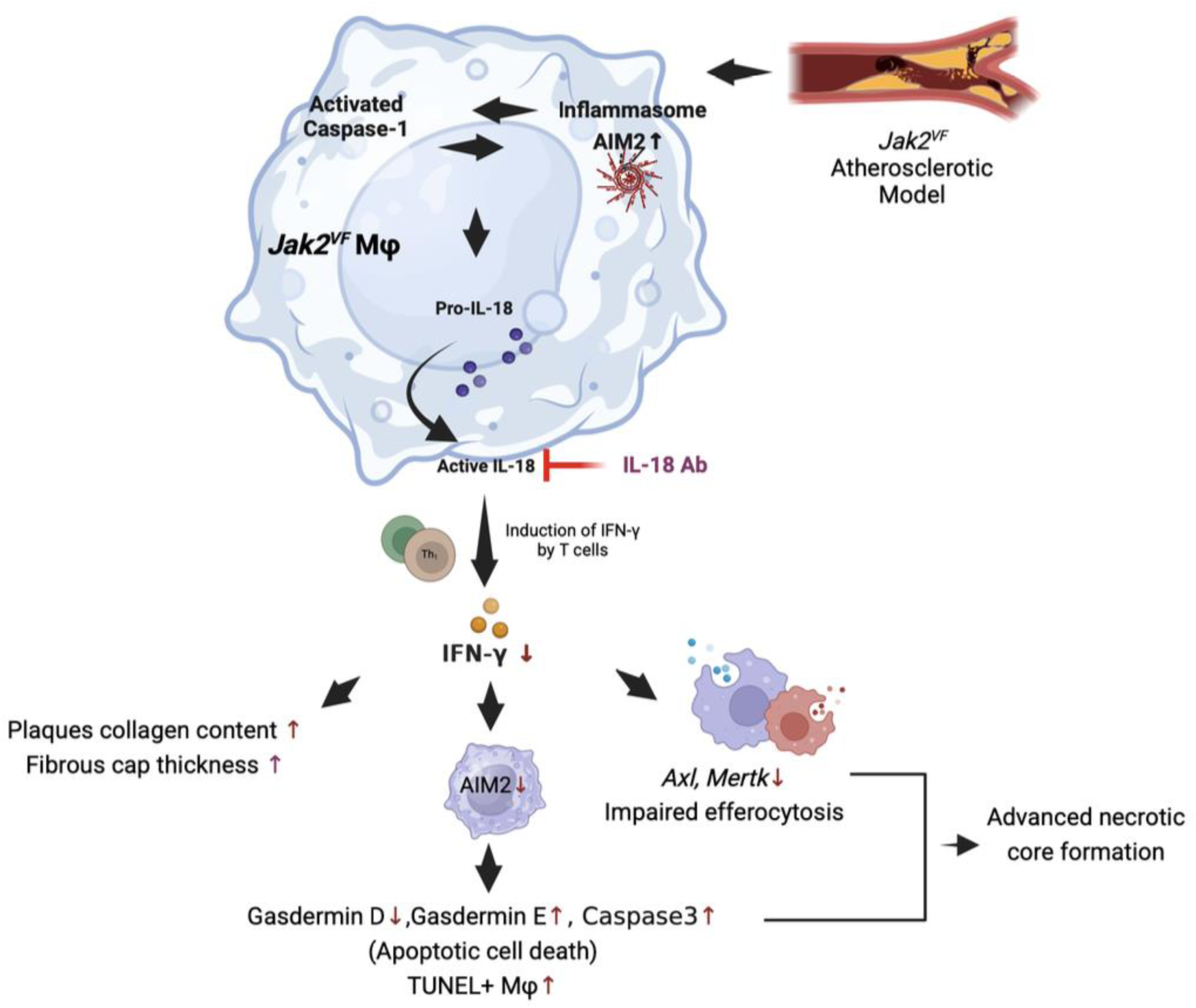

**Highlights:**

- Inflammasome activation produces active IL-1 and IL-18 and worsens atherosclerosis in clonal hematopoiesis (CH) however the contribution of IL-18 is unknown.
- Antibody inhibition of IL-18 increased plaque collagen but also increased early lesion area and late lesions with large necrotic cores in *Jak2^VF^* CH mice.
- There was a reversal of AIM2 inflammasome activation but a switch to apoptosis which along with reduced efferocytosis increased necrosis
- These events appeared to be coordinated by reduced IFN-γ which increased collagen but also decreased expression of efferocytotic genes. Our studies call into question whether inhibition of IL-18 would stabilize plaques in CH.

## Introduction

Clonal hematopoiesis (CH) arises from somatic mutations in hematopoietic stem cells (HSCs) that lead to the clonal expansion of blood cells, often involving leukocytes. CH increases with aging and augments overall mortality and cardiovascular risk. CH commonly involves the epigenetic modifiers *TET2*, *ASXL1*, *DNMT3A* or *JAK2^V617F^* (*JAK2^VF^*), that increases hematopoietic cytokine signaling. *JAK2^VF^* is less common than the other variants but arises at an earlier age and has the greatest impact on CVD risk. ^1, 2^ *Jak2^VF^* increases thrombosis and atherosclerosis in mouse models of CH and myeloproliferative neoplasm. ^3–5^ *Jak2^VF^* CH mutations lead to macrophage Absent in Melanoma 2 (AIM2) inflammasome activation and increased atherosclerosis in mice. ^5^ Inflammasome activation induces the secretion of active interleukin-1 beta (IL-1β) and interleukin-18 (IL-18). ^6, 7^ Elevated IL-18 levels are observed in both humans and mice with *JAK2^VF^* mutations. ^2, 4, 5, 8^ Although IL-1β antagonism in *Jak2^VF^*CH mice resulted in improved features of plaque stability, the role of IL-18 in atherogenesis in *Jak2^VF^* CH mice remains uncertain.

Previous studies have demonstrated heightened expression of IL-18 in unstable atherosclerotic lesions. ^9–11^ Studies in *Apoe^-/-^Il18^-/-^* mice revealed reduced lesion area and apparent plaque stabilization due to increased smooth muscle cell content. ^12^ IL-18 has been shown to promote atherosclerosis by stimulating interferon-gamma (IFN-γ) secretion from T cells, which in turn leads to fibrous cap thinning. ^10–15^ Consistently, IFN-γ increases plaque size and reduces plaque collagen content while IFN-γ receptor deficiency in *Apoe* knockout mice induces a marked increase in plaque collagen content, suggesting plaque stabilization. ^13–16^

While various studies in *Apoe* knockout or *Ldlr* knockout mice have implicated IL-1β in atherogenesis and features of plaque stability, 5^,17, 18^ the contribution of elevated plasma IL-18 to atherosclerosis in *Jak2^VF^* CH remains unclear. Increased AIM2 levels in *Jak2^VF^* CH, 5 could be driven by heightened IL-18-IFNγ signaling, contributing to AIM2 inflammasome activation and accelerated atherosclerosis. To investigate this hypothesis, IL-18 antibody therapy was employed in control and *Jak2^VF^*CH mice. IL-18 antibodies led to a reduction in IFN-γ and AIM2 levels and an increase in plaque collagen content, but unexpectedly also resulted in increased plaque area, necrotic core area and formation of advanced plaques with large necrotic cores. Further studies suggest an underlying mechanism involving a switch from pyroptotic to apoptotic cell death accompanied by a decrease in efferocytosis, calling into question the utility of IL-18 inhibition as a preventive measure for atherosclerotic CVD.

## Methods

### Study approval

All mouse experiments adhered to the guidelines of the Institutional Animal Care and Use Committee at Columbia University and were performed in accordance with the National Institutes of Health (NIH) Guide for the Care and Use of Laboratory Animals.

### 2.2. Animals

Mice used in these studies were female unless otherwise indicated and had a C57BL/6J genetic background. Mice with the *Jak2^V617F^* conditional knock-in allele were employed as described previously.5 Heterozygous *Jak2^V617F^*mice were crossed with heterozygous *Mx1-Cre* mice (B6. Cg-Tg (*Mx1-Cre*) (003356)) from Jackson Laboratories to produce *Mx1-Cre Jak2^VF^* mice. Additional animal colonies were acquired from the Jackson Laboratory, including wild-type C57BL/6J (000664), *Ldlr^−/−^*(B6.129S7-Ldlrtm1Her/J (002207)). Control mice included littermates carrying the Cre transgene.

### Bone marrow transplantation

Bone marrow transplantations (BMT) were carried out using established methods. ^5, 19^ Female *Ldlr^−/−^* recipient mice aged 8-12 weeks were lethally irradiated with a single 9.5 Gy dose from a cesium gamma and/or Multirad 350 source. Within 24 hours of irradiation, BM was collected from female donor mice aged 6-15 weeks with the specified genotypes. Donors were *Mx1-Cre Jak2^VF^* mice GFP^-^ (20%) and wild-type mice GFP^+^ (80%) to generate a *Jak2^VF^* clonal hematopoiesis (*Jak2^VF^* CH) model and corresponding controls. *Jak2^VF^*allele burden was assessed by flow cytometry as the percentage of GFP⁺ or GFP⁻ cells within the CD45⁺ population.

The irradiated mice were then randomized into treatment groups, anesthetized with isoflurane, and each received 3x10^6^ total bone marrow cells via intravenous (i.v.) injection in a final volume of 100 μl DMEM. Four weeks after BMT, cohorts with *Mx1-Cre* mice received intraperitoneal (i.p.) injections of 100 μg/mouse/day polyinosinic:polycytidylic acid (pIpC) twice, with a one-day interval between doses.

### Atherosclerosis studies and antibody administration

For atherosclerosis studies, *Ldlr^−/−^* recipient mice underwent BMT as described above and were subsequently fed a Western-type diet (WTD) (ENVIGO, TD.88137) for a specified duration beginning 6 weeks post-BMT. At 4 weeks after WTD feeding, mice were injected intraperitoneally with IL-18 antibodies (Bio X Cell, BE0237; 500 μg dissolved in 100 μl of in vivo dilution buffer (Bio X Cell, IP0070) or IgG isotype control (Bio X Cell, BE0089; 500 μg dissolved in 100 μl of in vivo dilution buffer (Bio X Cell, IP0070) three times per week with a 55-60 h interval between injections.

Following 8, 10, or 16 weeks of dietary intervention, mice were anesthetized with isoflurane at 5% in 3L/min of oxygen via inhalation for 5 minutes and blood was collected via the retro orbital vein for serum analysis, complete blood counts, and/or flow cytometric analysis. Following blood collection while remaining fully anesthetized as described above, mice were euthanized by cervical dislocation. Then hearts were harvested and fixed in 10% formalin solution for 48 hours. The fixed tissues were then embedded in paraffin and serially sectioned. Six sections which were evenly spaced, were stained with H&E to quantify total lesion and necrotic core areas. The reported necrotic area represents the average necrotic area per section. Additionally, cap thickness and collagen content was assessed after Masson trichrome staining (Sigma-Aldrich, HT15), following the manufacturer’s instructions.

Lesion size, necrotic core area, cap thickness, and collagen content were quantified using ImageJ (NIH) in a blinded fashion.

Atherosclerotic lesions were classified for severity, based on H&E staining, according to the American Heart System for humans, ^20^ which has been adapted for mice. ^21^ We based potential differences in lesion severity on the number of observations. We did 18 observations per mouse based on 6 H&E-stained sections per animal with 3 segments containing atherosclerotic lesions. The total observations per group is thus 18x the number of animals included in that particular group of mice.^22^

### Cap thickness measurement

For fibrous cap measurements, on the Masson trichrome (Polyscience, 25088-1) stained slides, we employed the method that was used in our previous studies. ^23^ We selected two slides on either side of the peak lesion area for evaluation of cap thickness. We captured images of the stained slides, including a scale bar, and analyzed using ImageJ software. For all observed lesions in a section, we measured the continuous collagen-stained region spanning the top of the lesion by grid lines. Then the average thickness of lesion was reported in length units.

### Complete blood counts

Complete blood counts were conducted on whole blood collected from retro orbital vein bleeding into an EDTA-coated tube. The samples were then analyzed with the Forcyte Veterinary Hematology Analyzer (Oxford Science).

### Serum analysis

Mouse serum was obtained by centrifuging blood at 12,000g for 10 minutes at 4°C. Serum IL-18 levels were measured using an ELISA kit (MBL International, 7625). Total serum cholesterol was assessed using a cholesterol E assay (Wako, 999-02601).

### Immunofluorescence Staining

Paraffin-embedded slides were first deparaffinized using Histo-clear and then rehydrated through a series of ethanol solutions with decreasing concentrations. Lesions were then stained with conjugated anti-mouse/human MAC2 AF488 (Galectin-3, Cedarlane, CL8942AF4, 1:100), anti-mouse Caspase-3 (Cell Signaling tech.,D175, 1:200), or anti-mouse Cl-GSDMD (R&D systems, CST 10137S, 1:10), anti-mouse GSDME-N-terminal (Abcam, Ab 215191, 1:100), anti-mouse AIM2 (Abcam, Ab119791, 1:200), anti-mouse IFN-γ (Invitrogen, MM700, 1:50) or anti-mouse AXL (R&D systems, AF854, 1:100), or rat isotype control IgG (BioLegend, 400431, 1:200), or rabbit isotype control IgG (Abcam, Ab172730, 1:150), and DAPI (BioLegend, 42280, 1:2500). Sections were incubated with primary antibodies at 4°C overnight, followed by a 30-minute incubation with secondary antibodies. For controls, isotype-matched IgG was used to ensure antibody specificity in immunofluorescence staining. TUNEL staining was conducted using commercial kit from Roche (12156792910). Images were captured using a green spinning-disk confocal microscope (Nikon Ti Eclipse inverted microscope).

### Flow cytometry

RBCs were lysed and samples were labeled with APC/Cy7 LIVE/DEAD (Invitrogen, MP34955) and incubated with PE anti-CD45 (BioLegend, 103106), PE-Cy7 anti-CD11b (BioLegend, 101216), BV421 anti-Gr-1 (BioLegend, 108445), and APC anti-CD115 (BioLegend, 134410), in staining buffer for 20 minutes at 4°C, and then washed once with staining buffer and resuspended in 10 μM H2DCFDA (Thermo Fisher Scientific, 88-5930-74) for another 15 minutes at 37°C. Samples were analyzed on a Novocyte Quanteon with Facs Software.

### Single-cell RNA Sequencing (Sc-RNA-seq) of Aortas from *Ldlr^−/−^* Mice

From the 16 weeks cohort on WTD, 6 mice from each group (*Ldlr^−/−^*mice transplanted with BM of *Jak2^VF^* CH mice or controls; treated with either IgG or IL-18 antibodies) were euthanized. The aortas were then carefully isolated and subjected to enzymatic digestion. The digestion process involved incubating the aortas with 2.5 mg/ml liberase (Millipore Sigma, LIBTM-RO), 120 U/ml hyaluronidase (Millipore Sigma, H3506), and 160 U/ml DNase I (Millipore Sigma, DN25) for 45 minutes at 37°C, following a previously established protocol. ^5^

Post-digestion, the cellular suspension was enriched for viable cells using a BD Influx sorter. To inhibit *ex vivo* necroptosis and gasdermin D oligomerization, 20 μM necro sulfonamide (Torcis) was added to all media used during the sorting process.

These sorted cells were then processed using the 10X Chromium single cell RNA seq platform (Single Cell 3′ Reagent Kits v2, 10X Genomics, USA) according to the manufacturer’s protocol.

### Sc-RNA-seq Data Analysis and batch effect correction

Aortic tissues from *Ldlr^-/-^* mice (Jackson, #002207) that had undergone bone marrow transplantation were analyzed using single-cell RNA sequencing (scRNA-seq). FASTQ files were processed using Cell Ranger v6.1.2, aligning reads to the mm10 mouse reference genome with Ensembl 93 annotation. Unique molecular identifier (UMI) count matrices were generated and filtered to retain cells expressing 200–2,500 genes, <10,000 UMIs, and <10% mitochondrial reads.

For dataset integration, previously published datasets GSE248395 24 and GSE163536 ^5^ were incorporated into the current study. Batch effect assessment was performed by comparing UMAPs before and after integration, revealing dataset-specific clustering that necessitated batch correction (Supplementary material online, Figure S1). 24 Canonical correlation analysis (CCA) with Seurat v5.0.3 was applied for integration. 24 Post-correction, samples clustered by cell type rather than dataset, confirming effective batch effect removal (Supplementary material online, Figure S1). ^5, 23, 25^

Downstream analysis included highly variable gene selection, normalization using log-transformed counts, and Louvain clustering (resolution = 0.5). Differential expression analysis was conducted using the Wilcoxon Rank Sum test, with Bonferroni correction (adjusted p < 0.05). Clusters were annotated based on known marker genes.

### Efferocytosis assessment and gene expression

To assess the effect of IFN-γ (PEPROTECH, 315-05) on gene expression and efferocytosis in *Jak2^VF^* CH and WT mice, male and female mice were divided into genotype and treatment groups (cytokines or vehicle). For gene expression analysis, mice received i.p. IFN-γ (50 ng/ml), and peritoneal macrophages were isolated 6 hours later using F4/80 Microbeads (Miltenyi Biotec). RNA was extracted (Qiagen, 74134) and analyzed by qPCR using SYBR Green (Applied biosystems, 4385612), normalizing to GAPDH.

For the efferocytosis assay, mice received i.p. IFN-γ, or PBS overnight. PKH26-labeled (Sigma-Aldrich, MIDI26) apoptotic Jurkat cells (1-2 × 10⁶) were injected ^26^, and peritoneal lavage was performed 2 hours later. F4/80+ macrophages were quantified for PKH26 positivity via flow cytometry as a measure of efferocytosis. ^27, 28^

### TUNEL assay

Samples were immune stained with TUNEL reagent (Roche), and anti-Mac2 antibodies (Cedarlane). Efferocytosis was measured by counting TUNEL+ cells that were associated with Mac2+ cells, that is, TUNEL+ nuclei in the cytoplasm of, or in contact with, Mac2+ macrophages, versus macrophage-free TUNEL+ cells, that is, TUNEL+ nuclei not in contact with neighboring macrophages.

### Statistical Analysis

Statistical analyses were performed using GraphPad Prism 10. The data are presented as mean ± standard error of the mean (s.e.m.). Measurements were obtained from distinct samples unless specified otherwise. Formal outliers were identified and removed using Grubb’s test.

For group comparisons, two-way analysis of variance (ANOVA) was employed. This provided information on treatment or genotype effects across groups and also assessed significance of genotype/treatment interactions. The results using this approach are shown in tables beneath the graphs. In addition, if significant ANOVA test results were obtained, we performed a post hoc Tukey’s multiple comparison test to identify which specific group means are significantly different from each other. The significant p values obtained by this approach are shown on the graphs.

An adjusted p-value <0.05 was considered significant and presented on the figures. The chi square test was used to determine differences in atherosclerotic lesion categorization. This test is based on the number of observations. ^22^ RNA-seq statistical analysis was conducted as previously described.^5, 23, 25^

## Results

*Ldlr^−/−^* mice were transplanted with a mixture of bone marrow from *Mx1-Cre Jak2^VF^*GFP^-^ mice (20%) and wild-type GFP^+^ mice (80%) to create a *Jak2^VF^* clonal hematopoiesis (CH) model, or with bone marrow from *Mx1-Cre* mice GFP^-^ (20%) and wild-type mice GFP^+^ (80%) as controls. Four weeks post-bone marrow transplantation (BMT), mice received pIpC injections to activate the *Jak2^VF^* mutation. Two weeks after the last injection, the mice were placed on a Western-type diet (WTD) for 8, 10, or 16 weeks to study atherosclerosis progression (Figure 1A-C). Starting at 4 weeks of WTD feeding, mice were treated with either isotype control IgG or IL-18 antibodies until the study’s conclusion (Figure 1A-C).

**Figure 1.**
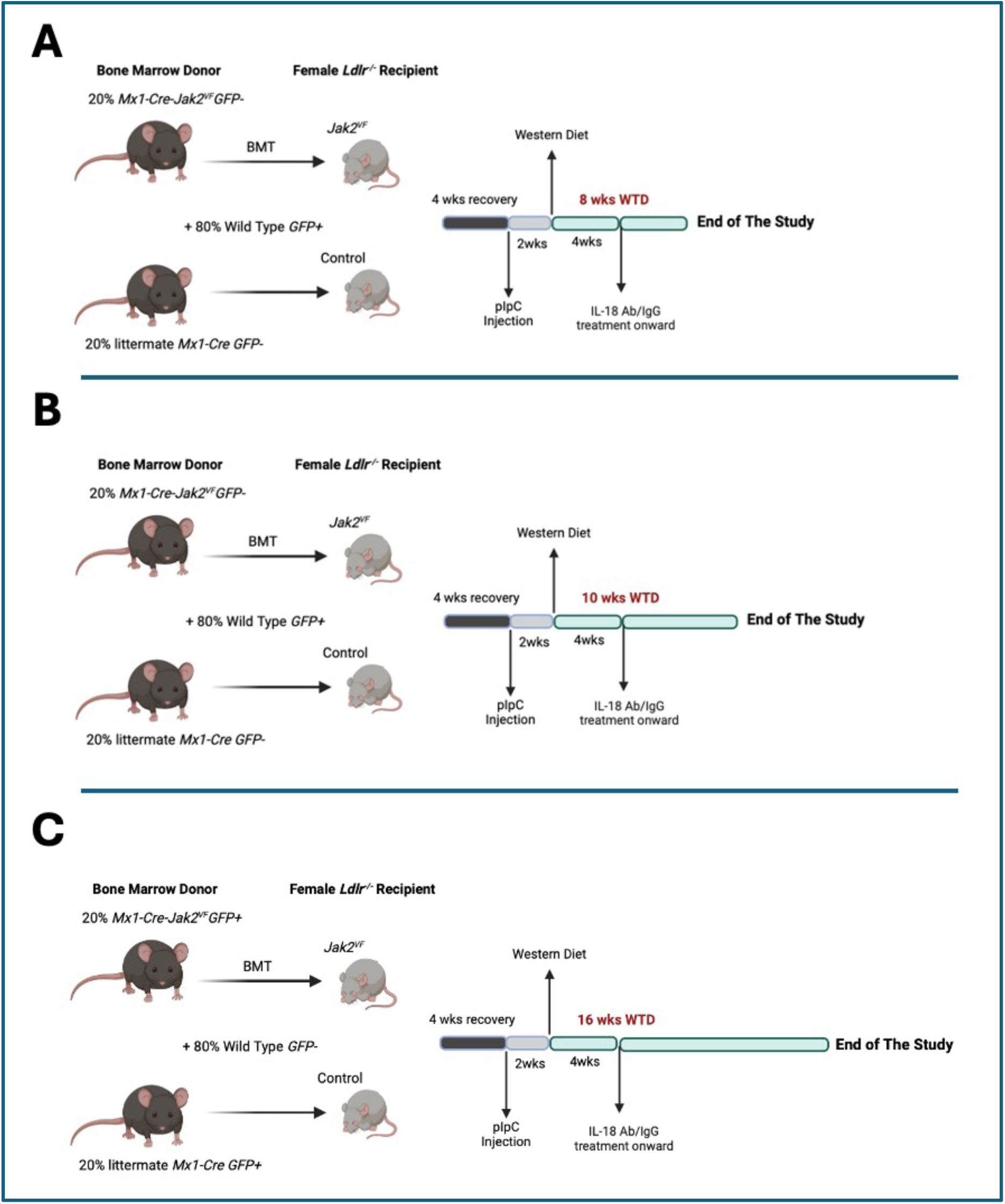
Experimental Design. *Ldlr^−/−^* mice were transplanted with a mixture of bone marrow (BM) from *Mx1-Cre-Jak2^VF^* (20%) and wild-type mice (80%) to generate a *Jak2^VF^* clonal hematopoiesis (CH) model or with BM from *Mx1-Cre* (20%) and wild-type mice as controls. At 4 weeks after BM transplantation (BMT), mice received two pIpC (100 µg per dose) injections to induce *Cre*. Two weeks later, mice were fed a Western-type diet (WTD) for a period of A. 8 (8 wks), B.10 (10 wks) or C. 16 weeks (16 wks). At 4 weeks of WTD feeding, mice were injected with IgG or Anti-IL18 for a period of 4 weeks, 6 weeks or 12 weeks respectively. The graphical illustrations were created with BioRender.com.

*Jak2^VF^* CH mice exhibited elevated blood leukocyte counts and hematocrit levels (Supplementary material online, Figure S2A-F). The mild increased *Jak2^VF^* allele burden in blood monocytes indicated successful engraftment and expansion of the mutant cells (Supplementary material online, Figure S2G) and was similar to earlier studies ^5^. Additionally, these mice showed decreased plasma cholesterol levels, larger spleens, and a reduction in overall body weight (Supplementary material online, Figure S3A-C). These findings are consistent with previous reports. **^Error! Reference source not found.^**, ^29–31^ None of these parameters were affected by IL-18 antibody treatment (Supplementary material online, Figure S3A-C). In a pilot study we found that IL-18 levels were elevated in *Jak2*^VF^ CH mice and significantly reduced by IL-18 antibodies (Supplementary material online, Figure S4A).

### Increased lesion area and advanced necrotic lesion formation in *Jak2^VF^*CH mice treated with IL-18 antibodies

Analysis of lesion area in the proximal aorta showed an overall increase with time on the WTD. Two-way ANOVA showed a significant increase in lesion area in *Jak2^VF^* CH mice by anti-IL-18 treatment at the 8-week time-point (Figure 2A-B) and a near significant treatment effect at 10-weeks (p=0.058, shown beneath the graph in Fig 2B). As expected, there was a significant genotype effect across groups at all time points. Overall, these findings indicate an adverse effect of IL-18 antibody treatment on lesion area in the *Jak2^VF^* CH mice, especially in early lesions.

**Figure 2.**
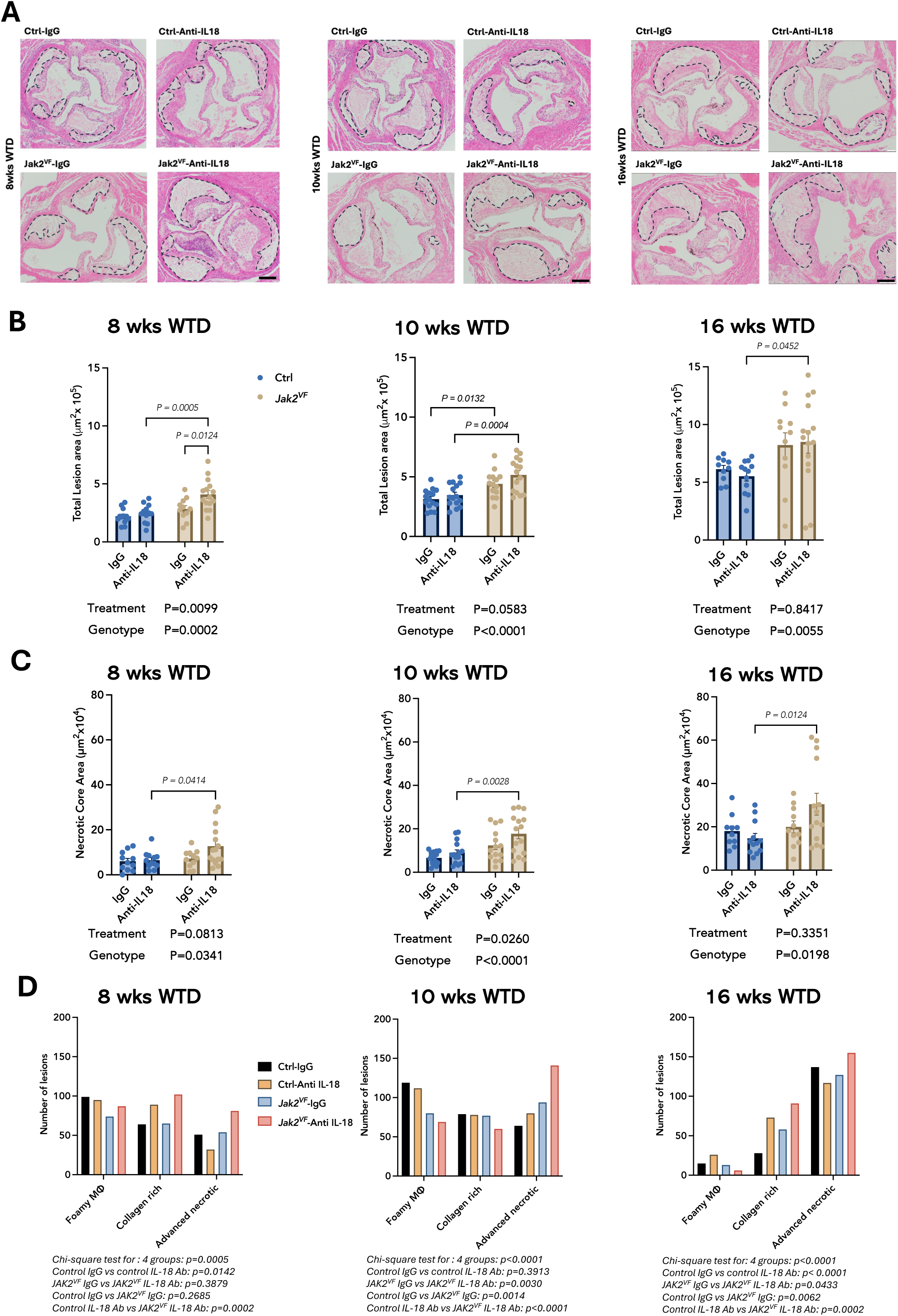
IL-18 antibody treatment increases lesion area and exacerbates necrotic core formation in *Jak2^VF^*CH mice. A. Representative hematoxylin and eosin (H&E) images of aortic root lesions. Necrotic cores are marked by dashed lines. Scale bar, 200 μm. B. Quantification of lesion area and C. necrotic core area. D. Number of foam cell rich lesions, collagen rich lesions, and advanced lesions with necrotic cores per group based on H&E staining and expressed per number of total observations. For B-D, n=12-15 mice for 8 wks, n=14 for 10 wks and n=10-14 for 16 wks cohort. For B and C, statistical analysis was performed using two-way ANOVA with Tukey’s multiple comparison test and for D using Chi-square test. Statistical differences are indicated on the graphs, with p-values greater than 0.05 omitted. Statistically significant differences by treatment and genotype factors are presented below the graphs.

We then examined features of advanced atherosclerosis, such as necrotic core area, a feature of plaque instability (Figure 2C). ^32–34^ Lesional necrotic core area was increased with time on the WTD. Analysis of data by two-way ANOVA showed a significant increase in necrotic core area following IL-18 antibody treatment at 10 weeks (treatment effect shown beneath the graphs in Figure 2A and C), as well as a genotype effect at all time points (shown beneath the graph). A further analysis of necrotic core area based on Masson trichrome staining showed a significant treatment effect of IL-18 antibody on necrotic core at 10 weeks and near significant treatment effects (p=0.05) at 16 weeks (Supplementary material online, Figure S4B). Genotype/treatment interactions were non-significant. These findings suggested that IL-18 antibody treatment increases necrotic core area at 10 week and possibly 16-week time points but did not allow us to conclude genotype specific effects.

To further assess the stage of lesion development, we employed a method that categorizes the type of lesions 21 among all groups by an experienced blinded observer, based on the number of observations per group. 22 This classification of plaques is categorical (non-parametric) and combines data based on all sections per plaque rather than averaging data obtained on a per mouse basis ^19, 22^. As expected, this showed a progression from foam cell rich lesions to advanced necrotic lesions with time on WTD. The analysis showed a significant increase in advanced lesions with large necrotic cores at 10 and 16-week time-points in *Jak2^VF^* CH mice treated with IL-18 antibodies (Fig 2D). In contrast, IL-18 antibody treatment did not have a major effect on large necrotic lesions at any time-point in control mice.

### IL-18 antibodies increase collagen content and fibrous cap thickness

To assess collagen content of lesions, we analyzed Masson Trichrome stained sections (Fig 3A). Consistent with earlier studies, ^9, 14^ there was a strong IL-18 antibody treatment effect on intimal collagen content at all time points (Fig 3A,B) The Tukey test showed a significant increase in collagen in response to IL-18 antibody treatment in controls at 8 weeks, and in *Jak2^VF^* mice at 10 and 16 weeks (Figure 3B). IL-18 antibodies significantly increased fibrous cap thickness at 8 and 10 weeks (treatment effect, Fig 3C).

**Figure 3.**
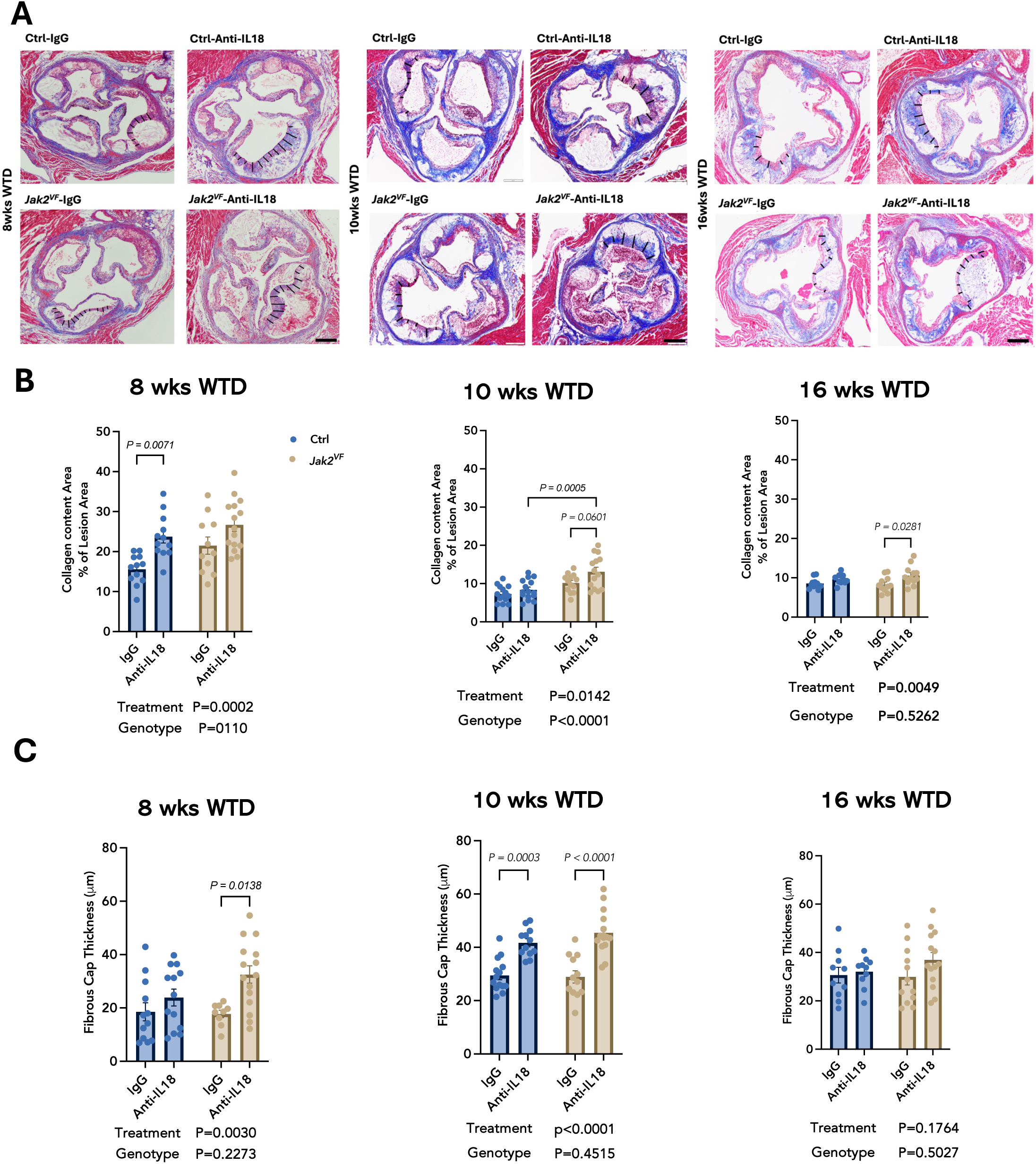
IL-18 antibody treatment enhances fibrous cap thickness and collagen content in atherosclerotic lesions. A. Representative images of Masson’s Trichrome staining showing collagen staining in lesions. Black bars indicate cap thickness. Scale bar, 200 μm. B. Total collagen staining (Blue) in lesions and C. Quantification of fibrous cap thickness (n=12-15 mice for 8 wks, n=14 for 10 wks and n=10-14 for 16 wks cohort) at 8, 10 and 16 weeks WTD feeding. For B and C statistical analysis was performed using two-way ANOVA with Tukey’s multiple comparison test. Statistical differences are indicated on the graphs, with p-values greater than 0.05 omitted. Statistically significant differences by treatment and genotype factors are presented below the graphs.

### Mechanisms of increased necrosis

While our findings on plaque collagen content are consistent with earlier studies, ^9, 14^ the increase in lesion area and necrotic core-rich lesions was unexpected. We next performed studies to assess potential mechanisms underlying increased necrotic core formation, focusing on 10- and 16-week WD feeding data showing increased advanced necrotic lesions. Our earlier studies indicated a role of AIM2 inflammasome activation and gasdermin D (GSDMD)-mediated pyroptosis in necrotic core formation in *Jak2^VF^*CH mice. ^5^ IFN-γ increased AIM2 expression in bone marrow-derived macrophages (BMDMs) and plaques from *Jak2^VF^*CH mice showed increased AIM2 levels. ^5^ Two way ANOVA showed that at 10 and 16 weeks WD feeding, IFN-γ levels were significantly reduced by IL-18 antibody treatment (Figure 4A-B, treatment effect). At 10 weeks there were significant increases in IFN-γ and significant reductions by treatment specifically in the *Jak2^VF^* group. IL-18 antibody treatment also reduced AIM2 levels at 10 and 16 weeks (Figure 4C-D, treatment effect), with a significant reduction by treatment in the *Jak2^VF^*group at 16 weeks. Moreover, an antibody that recognizes the cleaved, active form of GSDMD 35 showed significant reductions in cleaved GSDMD by anti-IL-18 treatment at 10- and 16-week time-points (Figure 4E-F, treatment effect). There were significant reductions in the elevated cleaved GSDMD levels at both time points in the *Jak2^VF^* groups. These findings suggest a complete reversal of increased AIM2 inflammasome activation by IL-18 antibody treatment, in parallel with a reduction in IFN-γ expression. Thus, the increase in advanced necrotic lesions in *Jak2^VF^* CH mice treated with IL-18 antibodies cannot be attributed to an increase in GSDMD-mediated pyroptosis, indicating an alternative mechanism.

**Figure 4.**
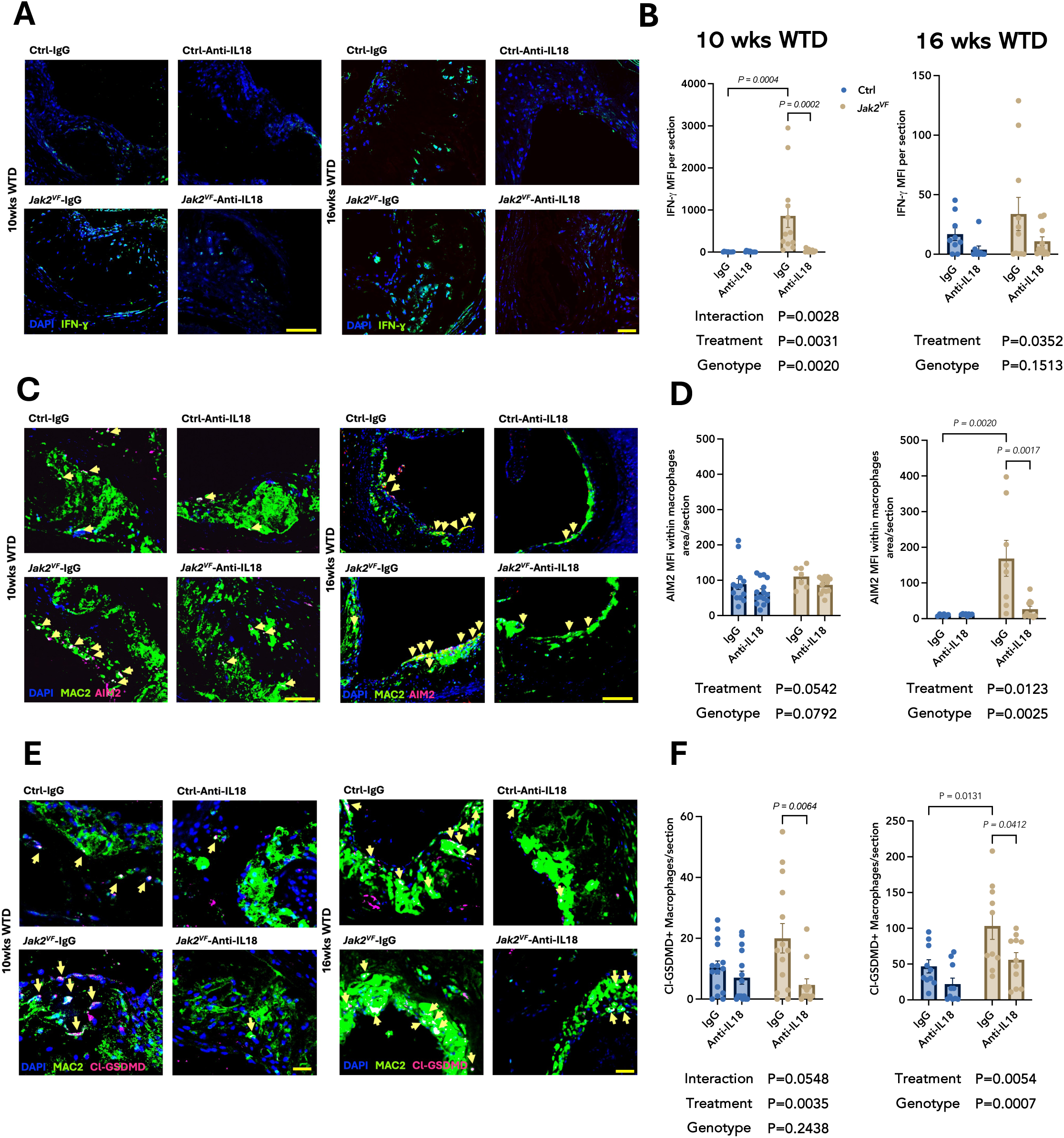
IL-18 antibody treatment decreased cleaved gasdermin D (Cl-GSDMD) levels, interferon (IFN)-γ and AIM2 expression in atherosclerotic lesions of *Jak2^VF^*CH mice. A-B. IFN-γ immunostaining. A. Representative images. IFN-γ is shown in green. B. Quantification (n=12-15 mice for 8 wks, n=12 for 10 wks and n=10 for 16 wks cohort). C-D. AIM2 inflammasome expression in macrophages in lesions. C. Representative images. AIM2 is shown in magenta. AIM2 positive macrophages (white spots) are highlighted by yellow arrows in lesions. D. Quantification (n=12 mice for 10 wks and n=8-14 for 16 wks cohort). E-F. Cl-GSDMD expression in macrophages in the lesions. E. Representative images. Cl-GSDMD is shown in magenta. Cl-GSDMD positive macrophages (white spots) highlighted by yellow arrows in lesions F. Quantification (n=15 mice for 10 wks and n=10 for 16 wks cohort). Scale bar for all images, 100 μm. For B, D, and F statistical analysis was performed using two-way ANOVA with Tukey’s multiple comparison test.

Macrophage apoptosis plays an important role in necrotic core formation in advanced necrotic lesions. 33^, 34, 36^ Thus, we next considered a potential role of apoptotic cell death, using cleaved caspase-3 and TUNEL staining as markers of apoptosis. Macrophage cleaved caspase-3 was increased by IL-18 antibodies at both time points (Figure 5A-B, treatment effect). IL-18 antibody treatment significantly increased TUNEL+DAPI-apoptotic corpses at 10 and 16 weeks (Figure 5C-D, treatment effect). There was no significant genotype/treatment interaction. At 16 weeks there was a significant increase in apoptosis by IL-18 antibodies in both groups. Together, these findings suggest that the increase in necrotic core formation with IL-18 inhibition might reflect a switch in the mode of cell death from GSDMD-mediated pyroptosis to post-apoptotic necrosis.

**Figure 5.**
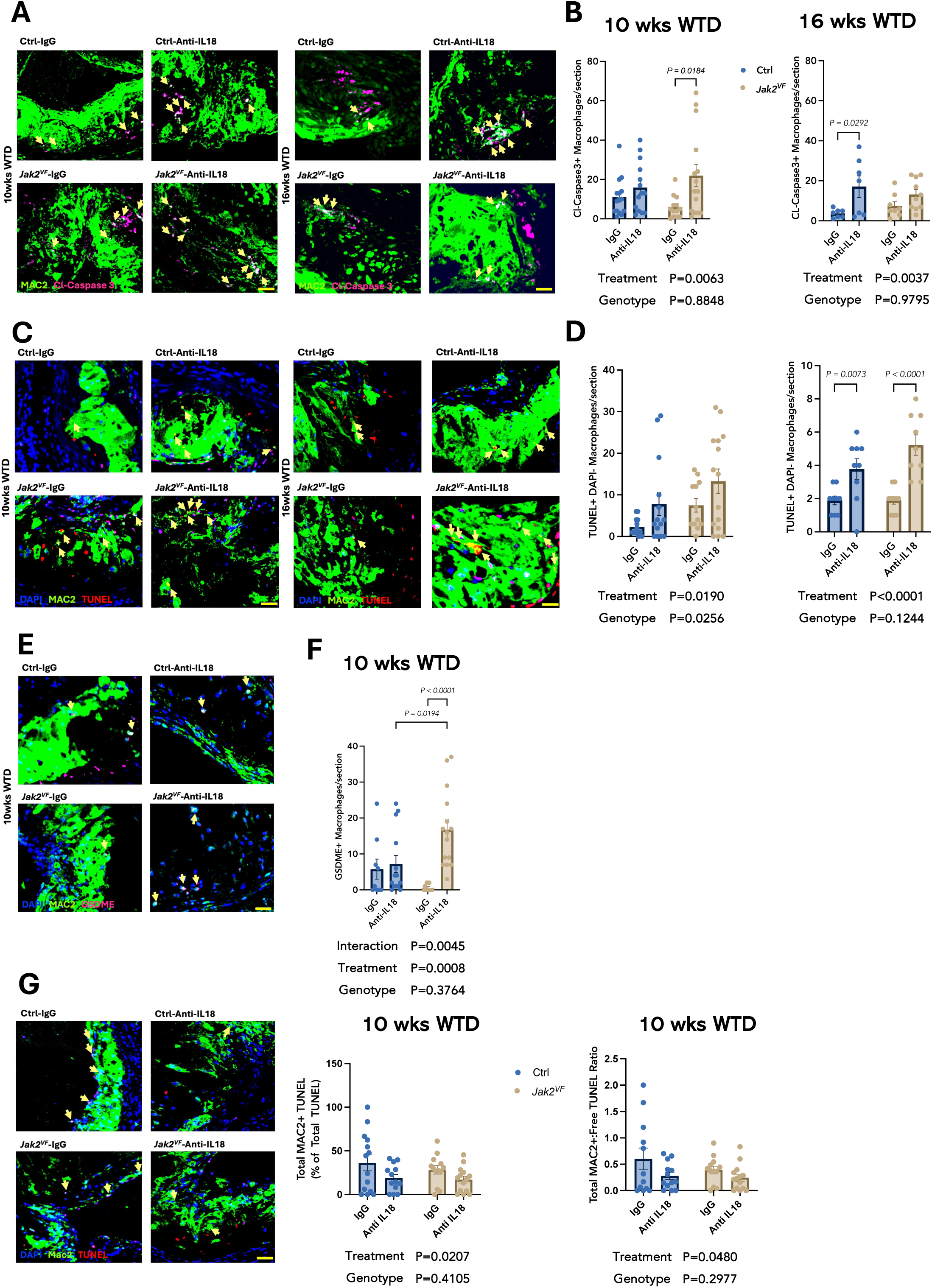
IL-18 antibody treatment increases apoptotic macrophages and impairs efferocytosis. A-B. Apoptotic cleaved (Cl)-Caspase-3+ macrophages. A. Representative images. Cl-caspase-3 is shown in magenta. Cl-caspase-3 positive macrophages (white spots) highlighted by yellow arrows in lesions. B. Quantification (n=13-15 mice for 10 wks and n=7-10 for 16 wks cohort). C-D. TUNEL assay. C. TUNEL^+^ staining visualized in red spots; dead macrophages (yellow spots) indicated by yellow arrows. D. Quantification (n=15 mice per group for 10 wks and n=10 for 16 wks cohort). E-F. GSDME, a pyroptotic marker, in macrophages. E. Representative images. GSDME is shown in magenta. GSDME positive macrophages (white spots) highlighted by yellow arrows in lesions.F. Quantification (n=15 mice per group for 10 wks and n=10 for 16 wks cohort). G. Representative images and in situ efferocytosis assessment (n=15 mice per group for 10 wks and n=10 for 16 wks cohort). Scale bar for all images, 100 μm. For B, D, and F statistical analysis was performed using two-way ANOVA with Tukey’s multiple comparison test.

While activated caspase-3 can inactivate GSDMD, ^37, 38^ it can also *activate* gasdermin E (GSDME), which in turn can promote post-apoptotic necrosis or pyropotosis. ^38–40^ Consistent with this scenario, GSDME levels were increased in *Jak2^VF^* CH mice by IL-18 antibodies at 10 weeks (Figure 5E-F, treatment effect). Similar results were obtained using an antibody that recognizes the N-terminal active fragment of GSDME (p=0.05 for treatment effect) (Supplementary material online, Figure S5A-B). These findings suggest that an increase in GSDME mediated apoptosis may increase necrotic core formation following IL-18 antibody treatment. To assess the possibility that there might also be a defect in efferocytosis, we determined the efficiency of efferocytosis, expressed as the % macrophage-associated to total TUNEL+ bodies or the ratio macrophage-associated to free TUNEL+ bodies. 41 This revealed a decrease in the efficiency of efferocytosis at the 10-week time-point, as suggested by a significant treatment effect (Figure 5G).

### Single Cell RNA-seq Analysis

To gain further insights into the mechanisms responsible for the worsening of plaque features in *Jak2^VF^* mice treated with IL-18 antibodies, we integrated scRNAseq datasets of aortic CD45^+^ cells isolated from *Jak2^VF^* CH mice with IL-18 or IgG antibodies and datasets from previous studies.^5, 23^ The integrated analysis of mouse aortic leukocytes allowed us to identify 17 clusters from 11,838 cells (Figure 6A). Consistent with prior reports **^Error! Reference source not found.^***Jak2^VF^* CH mice displayed an increased fraction of lesional resident-like macrophages (cluster 0, Figure 6B) and a decreased fraction of lesional Trem2-like macrophages (cluster 2, Figure 6B). However, there was no marked alteration in the population distribution of lesional cells resulting from IL-18 antibody treatment (Figure 6B). Since defective efferocytosis can contribute to increased necrotic core formation, ^23, 42^ we assessed the expression of efferocytosis genes in macrophage subpopulations. Anti-IL-18 treatment decreased *Axl*, *Mertk*, and *Cd36* expression in resident macrophages and *Axl* expression in one population of Trem2+ macrophages (Figure 6C). Immunostaining suggested a decrease in macrophage AXL protein expression in both groups of mice in response to IL-18 antibody treatment with a significant treatment effect shown by two-way ANOVA (Figure 6D-E). These findings suggest that a decrease in expression of efferocytosis genes might contribute to decreased efferocytosis in *Jak2^VF^* mice treated with IL-18 antibodies.

**Figure 6.**
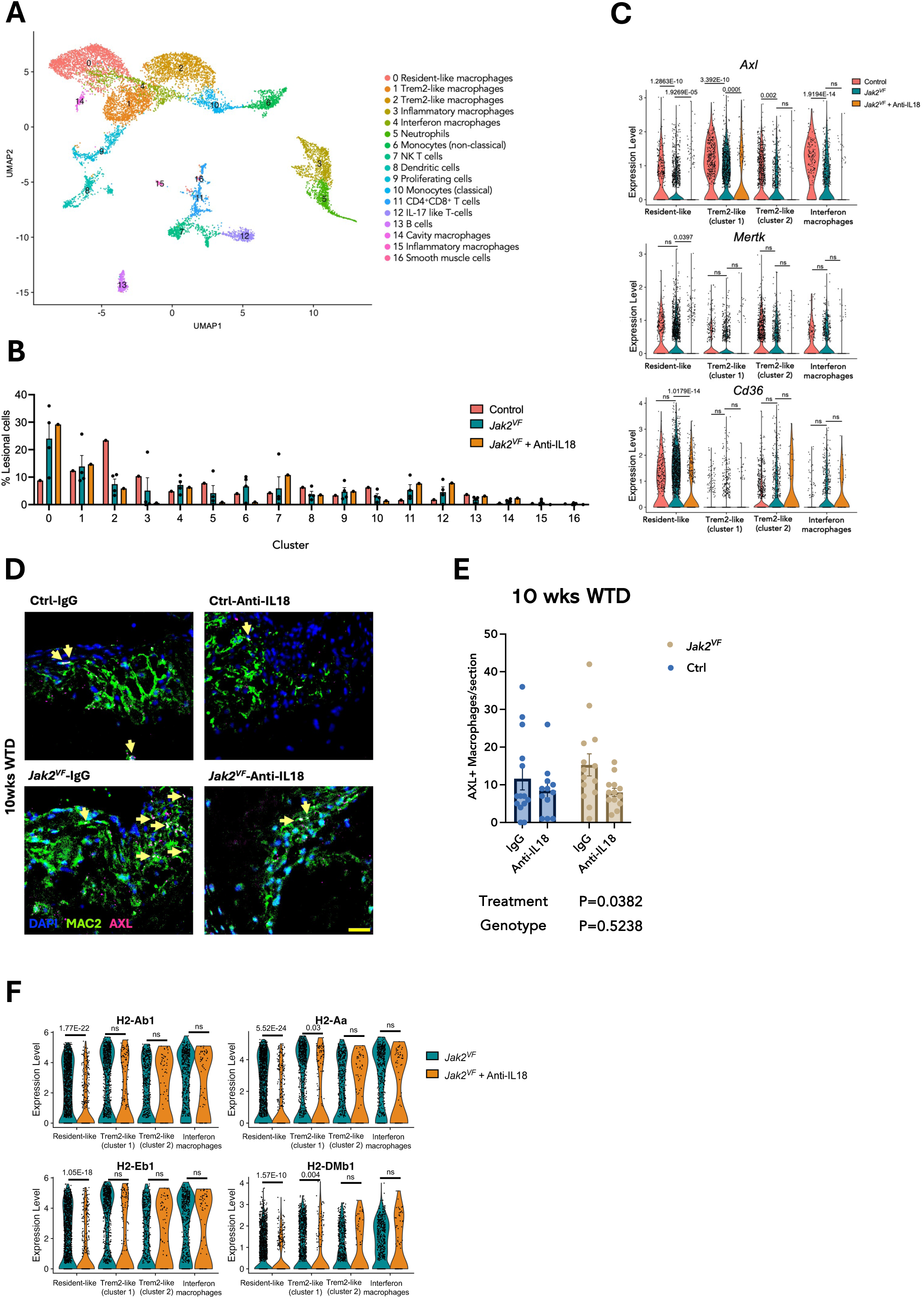
Singe cell RNA-seq analysis of atherosclerotic plaques shows reduced expression of efferocytosis genes in resident-like macrophages. A. UMAP of integrated single-cell RNA-sequencing of CD45+ plaque cells isolated from aortic arches of control mice, *Jak2^VF^* mice, and *Jak2^VF^* mice with IL-18 antibodies. B. Percentage of lesional CD45^+^ cells within each cluster. C. Violin plots depicting gene expression levels of efferocytosis markers *Axl*, *Mertk*, and *Cd36* in control, *Jak2^VF^*, and *Jak2^VF^* mice treated with IL-18 antibody, derived from Sc-RNA-seq data. D. Representative aortic images with DAPI, MAC2, and AXL staining. DAPI is shown in blue, MAC2 is shown in green, and AXL is shown in magenta. AXL positive macrophages (white spots) are highlighted by yellow arrows in the lesions. E. Quantification of lesional AXL positive macrophages (n=15/group) after 10 weeks of WTD feeding. F. Violin plots depicting gene expression levels of MHC II genes in *Jak2^VF^*, and *Jak2^VF^* mice treated with IL-18 antibody, derived from Sc-RNA-seq data. Scale bar, 100 μm. Statistical analyses were performed using the Wilcoxon Rank Sum test with Bonferroni correction in C. For E statistical analysis was performed using two-way ANOVA with Tukey’s multiple comparison test. Statistical differences are indicated on the graphs, with p-values greater than 0.05 omitted. Statistically significant differences by treatment and genotype factors are presented below the graphs.

### Mechanisms of reduced efferocytic gene expression

Further studies were performed to assess the mechanisms of reduced expression of efferocytic genes. Prior studies have suggested that IFN-γ treatment could increase macrophage efferocytosis. ^28^ Analysis of Sc-RNA-seq data showed a marked decrease in mRNA expression of *MHC II* genes in macrophages (Figure 6F), strongly suggesting a decrease in IFN-γ signaling. ^43^ This decrease was restricted to resident-like macrophages and one population of Trem2-like macrophages, that also displayed the most pronounced changes in expression of efferocytosis genes (Fig 6C). To directly test a possible role of IFN-γ in inducing efferocytic genes we injected WT or *Jak2^VF^*mice with IFN-γ or control lgG and then isolated resident peritoneal cavity macrophages (Figure 7A). There was a significant increase in *Mertk* and *Axl* mRNA in *Jak2^VF^*female mice, with a trend upwards in controls (Figure 7B). In similar experiments *Axl* but not *Mertk* mRNA was increased by IFN-γ in male mice suggesting a possible sex difference in the response (Figure 7C). An assessment of efferocytosis using flow cytometry ^41^ showed a significant increase in IFN-γ treated *Jak2^VF^* female mice. These results suggest that the decrease in IFN-γ levels in plaques and in IFN-γ signaling in macrophage subpopulations observed following IL-18 antibody injection might be responsible for the reduced expression of efferocytosis genes contributing to a decrease in efferocytosis in resident-like macrophages.

**Figure 7.**
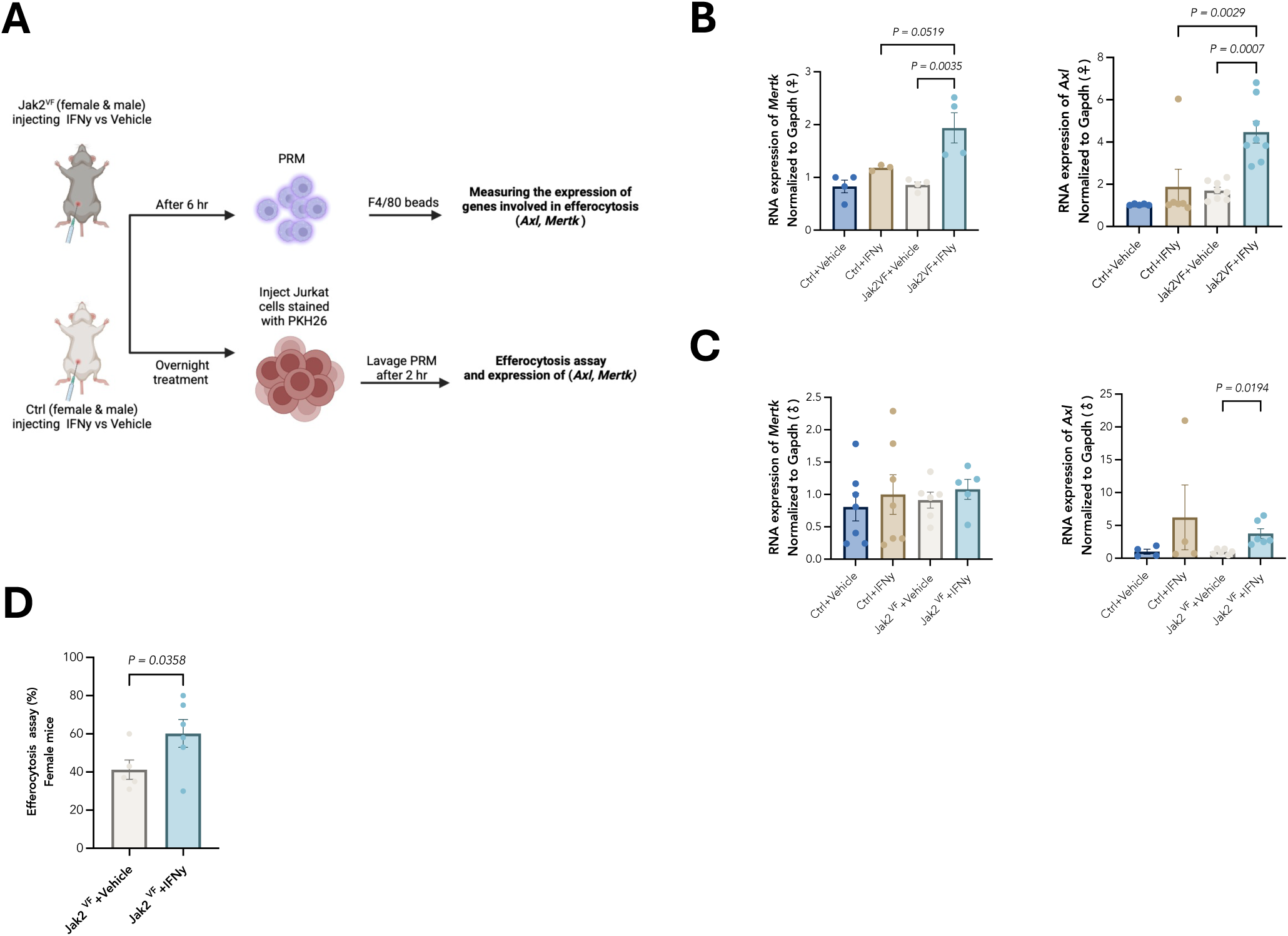
IFN-γ treatment enhances efferocytosis and expression of efferocytosis genes in resident peritoneal macrophages. A. Experimental setup. *Jak2^VF^* mice and their littermate controls (males and females) were injected with IFN-γ. B-C. At 6 hr after injection, resident peritoneal macrophages were isolated and mRNA expression levels of efferocytosis markers *Axl* and *Mertk* were assessed in Female (B) or male (C) (n=3-6/group). D. Overnight after the initial injection, mice were injected with PKH26 stained Jurkat cells, and two hours later the phagocytosis of Jurkat cells by resident peritoneal macrophages was assessed by flowcytometry(n=5-6/group). For B and C statistical analysis was performed using ordinary one-way ANOVA with Tukey’s multiple comparison test. For D, t-test was applied.

## Discussion

Previous studies have demonstrated the role of inflammasomes in promoting atherosclerosis and features of plaque destabilization in *Jak2^VF^* and *Tet2* CH mice. ^5, 7, 44–46^ Inflammasome activation triggers the secretion of both active IL-1β and active IL-18, and IL-18 levels are prominently increased in plasma of *Jak2^VF^* mice and humans.^2, 8^ IL-18 has been implicated in promoting atherosclerosis through IFN-γ-mediated inflammatory amplification and plaque destabilization ^10, 13, 14^. In the present study, treating atherosclerotic *Jak2^VF^* CH and control mice with IL-18 antibodies resulted in decreased plaque IFN-γ levels, thickened fibrous caps, and increased plaque collagen content, in line with prior findings demonstrating the role of IFN-γ in reducing collagen content of plaques. ^14, 15^ However, IL-18 antibody treatment also increased early lesion area and increased intermediate and advanced lesions with large necrotic cores in *Jak2^VF^* CH mice. Our studies suggest a previously unsuspected adverse effect of IL-18 antagonism on early lesions and on necrotic core formation, which is considered a key feature of plaque instability. ^32–34^ We found that IL-18 blockade led to decreased levels of IFN-γ and reduced expression of AIM2, which is known to be induced by IFN-γ. ^5, 47, 48^ In prior studies we have shown that AIM2 inflammasome suppression and IL-1β inhibition increased fibrous cap thickness in *Jak2^VF^* CH mice. ^23^ The observed reduction in AIM2 and IFN-γ levels following IL-18 neutralization thus represent a plausible mechanism to explain the increase in plaque collagen content.

Prior studies of IL-18 or IFN-γ suppression did not specifically report on the necrotic core area of atherosclerotic plaques. ^12–15^ Assessing multiple time points of WTD feeding, we found a significant increase in advanced lesion sections characterized by very large necrotic cores at 10 and 16 weeks in *Jak2^VF^* but not control mice. Necrotic core area showed a significant or near significant IL-18 neutralization effect at 10 and 16 weeks WTD feeding (treatment effect by two-way ANOVA) with similar trends in controls and Jak2VF mice. An increase in cleaved caspase-3 and apoptotic macrophages was similarly observed in both control and *Jak2^VF^* mice. However, in the 10 week data set, the increase in IFN-γ and its reduction by treatment, as well as the decrease in pyroptosis and the increase in GSDME were specific for the *Jak2^VF^*mice. Together, these findings suggest a common underlying mechanism of plaque necrosis in response to IL-18 inhibition that may become more prominent in a setting where IL-18 production is increased such as Jak2VF CH. Further assessments of IL-18 inhibition in other settings where IL-18 levels are increased such as obesity and diabetes may be warranted. ^49–51^

Seeking to understand the mechanism of increased necrotic core formation, we utilized a Cl-GSDMD antibody, which specifically recognizes the active amino-terminal (N-terminal) fragment. ^35^ This revealed a decrease in Cl-GSDMD, paralleling the decrease in AIM2 expression. In contrast, TUNEL^+^ and cleaved Caspase-3^+^ macrophages were markedly increased by IL-18 antibody treatment, indicating an increase in apoptotic cell death. This suggests that IL-18 blockade causes a switch from pyroptosis to secondary apoptosis. Alternation between different modes of cell death has been described in other settings. ^38, 52^

Active caspase-3 has been reported to disable GSDMD by cleaving its N-terminal pore-forming domain at a specific site, ^38, 53, 54^ which along with the decrease in AIM2 expression may explain the decrease of Cl-GSDMD due to IL-18 antibody treatment. Furthermore, the proteolytic activation of GSDME by caspase-3 can induce either pyroptosis or secondary membrane damage after apoptosis. ^53–55^ Prior studies have indicated reduced *Mertk* and defective efferocytosis in *Jak2^VF^* mice.^56, 57^ Decreased expression of *Mertk*, *Axl* and *Cd36* in resident-like macrophages in response to IL-18 antibody treatment suggests a treatment-related impairment of efferocytosis. The combined effects of increased apoptosis and decreased efferocytosis explain the increase in plaque necrosis in *Jak2^VF^* mice treated with IL-18 antibodies.

A key observation by scRNA-seq was that gene expression of MHC II subunits was decreased in resident-like macrophages in *Jak2^VF^* CH mice treated with IL-18 antibodies, suggesting a decrease in IFN-γ signaling and correlating with decreased expression of efferocytosis genes in the same macrophage subpopulation. Prior studies have shown that interferon-γ increases macrophage efferocytosis 28 and we showed that IFN-γ treatment increased expression of *Mertk* and *Axl* suggesting induction of efferocytosis genes as an underlying mechanism. The larger effect in *Jak2^VF^* compared to WT macrophages could indicate increased IFN-γ signaling as a result of the *Jak2^VF^*mutation and could explain the more pronounced effects of IL-18 inhibition on lesional phenotypes in *Jak2^VF^* versus control mice.

IL-18 stimulates IFN-γ production by T cells in plaques. ^12^ In our study the different effects of IL-18 inhibition on plaque phenotype may be coordinated by the decrease in IFN-γ, which increases plaque collagen and fibrous cap thickness, reduces AIM2 expression and inflammasome activation, activating a switch to Caspase-3 activation and apoptosis, and reduces expression of efferocytosis genes in macrophages. This leads to accumulation of apoptotic cells and post-apoptotic necrosis (Graphical abstract). More broadly our findings emphasize the offsetting effects of IFN-γ signaling on features of atherosclerotic plaque stability.

Targeting inflammasomes in inflammatory disorders is an active and promising area of investigation. 58^, 59^ For example, two compounds DFV890, an NLRP3 inhibitor and MAS825 a bispecific antibody that target IL-18 and IL-1β, are novel investigational drugs that are being assessed for reduction of inflammatory biomarkers in CH patients with atherosclerotic cardiovascular disease (ACVD) (*Clinical Trials.gov; #NCT06097663*). Our studies in a preclinical model suggest that IL-18 neutralization, by giving rise to mixed atherosclerosis phenotypes, may not to be an optimal choice for the treatment of atherosclerosis associated with *Jak2^VF^* CH, or perhaps more generally in other conditions where elevated IL-18 levels or production result from inflammasome activation. Consistent with our findings of a mixed atherosclerosis phenotypes, genetically predicted reductions in IL-18 levels in humans reduced IFN-γ levels but did not alter the risk of coronary artery disease or ischemic stroke suggesting no benefit for atherosclerotic disease. ^60^ In contrast, IL-1β inhibition which improves features of plaque instability ^61^ and decreases ACVD in humans ^62^ could be a preferable approach for reducing AVCD in selected individuals with CH, offering targeted therapy without compromising the delicate balance of cytokine signaling and cell death mechanisms in atherogenesis. ^23, 63, 64^

## Acknowledgements

This study used the Confocal and Specialized Microscopy Shared Resource of the Herbert Irving Comprehensive Cancer Center at Columbia University, funded in part through NIH/NCI Cancer Center Support Grant P30CA013696. The authors thank Dr. Sekhar Ramakrishnan (Department of Pediatrics, Columbia University) for statistical advice.

## Author contribution statement

M.T., A.T., and M.W. made substantial contributions to the conception, design, data acquisition, analysis, and interpretation. They also drafted and critically reviewed the manuscript for intellectual content. C.H. and B.H. contributed to data analysis. The remaining authors reviewed the manuscript and approved the final version for publication.

## Non-Standard Abbreviations and Acronyms

*Jak2^V617F^*: Januse Kinase 2 ^V617F^
IL-1β: Interlukin 1-Beta
IL-18: Interlukin-18
CH: Clonal Hematopoiesis
BM: Bone Marrow
WTD: Western Type Diet
IFN-γ: Interferon Gamma
AIM2: Absent in Melanoma 2
TUNEL: Terminal deoxynucleotidyl transferase mediated d-UTP nick end labeling
Sc-RNA-seq: Single cell RNA Sequensing
MHC II: Major Histocompatibility Complex 2
*Mertk*: Myeloid Epithelial Tyrosine Kinase
*Axl*: AXL receptor tyrosine kinase
HSCs: Hematopoietic Stem Cells
*TET2*: Tet methylcytosine dioxygenase 2
*ASXL1*: Additional sex combs like 1
*DNMT3A*: DNA Methyltransferase 3 Alpha
CVD: Cardiovascular Disease
Apoe: Apolipoprotein
E GSDMD: Gasdermin D
GSDME: Gasdermin E

